# Mapping the Cerebrovascular Abnormality in Transgenic Alzheimer’s Disease (AD) Mice with deep-learning-based super-resolution cerebral blood volume (CBV)-MRI

**DOI:** 10.1101/2024.05.17.594785

**Authors:** Xiaoqing Alice Zhou, Xiaochen Liu, Hongwei Bran Li, David Hike, Yuanyuan Jiang, Matthew S. Rosen, Juan Eugenio Iglesias, Xin Yu

**Author notes:** Address correspondence to: Xin Yu, Ph.D. These authors contribute equally to the paper (Xiaoqing Alice Zhou, Xiaochen Liu).

## Abstract

To measure the brain-wide vascular density (VD) alteration in degenerated brains with Alzheimer’s Disease (AD), deep learning-based super-resolution approach was developed to assist the segmentation of micro-vessels from the Monocrystalline Iron Oxide Nanoparticle (MION)-based CBV MRI images of transgenic mouse brains. Iron-induced T2* amplification effect well separated micro-vessels with tens of microns from capillary-enriched parenchyma voxels, enabling vascular compartment-specific VD differential analysis between AD and wildtype control mice. The differential maps based on segmented micro-vessels identified decreased VD in the anterior cingulate cortex (ACC) and medial entorhinal cortex (mEC) and increased VD in several highlighted brain regions, including dentate gyrus (DG) of the hippocampus, central and geniculate thalamus, medial septal area (MS), ventral tegmental area (VTA), and lateral entorhinal cortex (lEC). In contrast, the T2*-weighted capillary density mapping from parenchyma voxels showed increased VD in several cortical regions, including somatosensory and visual cortex, retrosplenial cortex, as well as piriform area and lEC in AD brains. However, dramatic capillary VD decrease was observed in the subcortical areas including hippocampus, thalamus, hypothalamus, and pontine areas. These high-resolution MION-based CBV MRI elucidates altered vascular compartments in degenerated AD brains, reconciling the various region-specific vascular impairment and angiogenesis in functional areas critical for cognitive decline of AD.

## Introduction

Vascular abnormalities are crucial indicators of brain degeneration in aging or degenerative diseases, such as Alzheimer’s disease (AD). Early-stage AD exhibits impaired cerebrovascular function, marked by atrophy, diameter reduction, and other manifestations^1^. This vascular perturbation is theorized to be closely related to cerebral amyloidosis^2,3^ and Tau^4^, the hallmarks of AD, suggesting altered vascular dynamics as a key feature in AD pathology^5^. However, inconsistencies in the alterations of vascular density (VD)^6^, particularly with region specificity and disease progression dependency ^7^ suggest high variability of vascular pathology^8^. Extensive optical imaging or histological studies on transgenic rodent AD models report the reduced VD across different brain stages^3,9-18^. Whereas, two recent studies of transgenic mouse models have reported increased hippocampal vascular density at the early stage with a subsequent decline along disease progression^19,20^. In parallel, several AD human post-mortem studies revealed increased vascular density^21-24^, while many studies found either no significant change or a reduction in VD^25-44^. Although studies have reported the cerebral amyloid angiopathy (CAA), marked as the deposition of Aβ in the walls of blood vessels^45,46^, its brain-wide impact on cerebrovasculature in AD brains remains to be elucidated thoroughly either in human or transgenic animal models.

The conventional Magnetic Resonance Angiography (MRA) utilized in human and animal brain mapping has limitations in achieving the necessary resolution for micro-vessels with tens of micron diameter^47^. In contrast, emerging vascular mapping methods, including the doppler-based or micro-bubble ultrasound^48,49^, optoacoustic imaging^50,51^, and micro-CT^52-54^, have shown considerable advancements, in particular, offering ultra-high resolution cerebrovascular mapping in rodent brains. Despite the superior spatial resolution attained through either in vivo or ex-vivo brain sample, challenges persist. Instances of missed vessel detection arise due to potential vessel orientation issues, ineffective signal transmission from deep brain regions, and difficulties with resin filling through the cerebrovasculature for CT contrasts. There is still missing an effective *in vivo* method to map the brain-wide cerebrovasculature, meanwhile, holding the translational potential for non-invasive VD mapping of AD patients.

Magnetic iron oxide nanoparticles (MION)-based cerebral blood volume (CBV) mapping methods have been employed for *in vivo* brain-wide vasculatures^55^. The spatial resolution of MION-based CBV fMRI is lower than the aforementioned methodology. To achieve the voxel size with tens of micron resolution, elongated acquisition time is needed to produce the image with sufficient signal-to-noise ratio (SNR) when characterizing the cerebrovasulature with signal drop due to the iron particle-induced T_2_* decay. Since the SNR is increased by a factor of the square root of two given the increased acquisition time, existing studies faces the challenges to perform the ultra-high resolution MION-based CBV MRI for *in vivo* cerebrovascular mapping. Deep learning has emerged as a transformative technology to produce the super-resolution^56-59^ of MRI images based on the well-trained neural networks from existing cohort of datasets. By applying a novel deep learning-based super-resolution approach, it is feasible to improve the spatial resolution of MION-based MRI images, enabling brain-wide characterization of the micro-vessels, which are amplified spatially in the T_2_*-weighted CBV images.

In this study, we improved the MION-based CBV methods by incorporating a 3D deep learning approach, resulting in a finer reconstruction for micro-vasculature segmentation at a 38-micron cubic resolution from a 75μm cubic resolution raw images acquired in 24 min per animal using a 14T scanner. By employing a threshold-varied masking method, micro-vessels at different sizes are segmented into compartments, enabling the size-dependent density comparison. Meanwhile, by segmenting out the identifiable vasculature in MION-based CBV images, the signal intensity of the parenchyma voxels can be used to estimate the density of capillaries based on the MION-induced signal drop This newly developed mapping scheme enables a large cohort sample acquisition from both wildtype and transgenic mouse lines with Aβ overexpression. Voxel-wise statistical analysis of segmented vascular compartments and capillary density in parenchyma voxels enables the characterization of size-dependent vascular abnormalities associated with AD brain degeneration.

## Methods

### Animals

All animal surgical and experimental procedures were conducted following the Guide for the Care and Use of Laboratory Animals and approved by the Massachusetts General Hospital Subcommittee on Research Animal Care. Mice were kept on a 12/12-hour light cycle with unlimited access to food and water. 12 female APPswe/PSEN1dE9 (C57BL6) mice and 11 female C57BL/6 mice aged 6 - 10 months (25-35 g) were used for experiments.

### Surgical Procedures

Mice were subjected to a surgical procedure for the secure attachment of the RF coil to their heads. They were anesthetized with isoflurane, initiated at 5% concentration with a flow of 1L/min of medical air supplemented by 0.2L/min of O_2_. Anesthesia depth was carefully monitored by observing the animals’ respiration rates, maintaining isoflurane levels at 1.5%-2%. The heads were immobilized on a stereotaxic apparatus using ear and bite bars. After shaving and sterilizing the scalp with ethanol and iodine, an incision was made to prepare a skull area matching the RF coil’s ring size. The skull was cleared of any tissue remnants, treated with 0.3% hydrogen peroxide and phosphate-buffered saline, and dried. The coil was carefully aligned and placed on the skull, elevated approximately 0.3-0.5mm to prevent excessive pressure and secured in place using a thin layer of cyanoacrylate adhesive. After the adhesive set (∼5-8 minutes), dental cement was mixed and applied to encapsulate the coil and the skull, ensuring the base was solidly anchored and being cautious to prevent any cement from dripping towards the eyes. The incision was then sealed. Post-operation, the mice received subcutaneous injections of Buprenorphine and Cefazolin and were returned to their home cage for a minimum of one-week recovery, allowing for adequate neck muscle strengthening for normal head movement. For in vivo vasculature mapping, mice were injected with 2.5mg/kg of MION particles from Biopal (10mg/ml solution) via tail vein half an hour before MRI scanning.

### Magnetic Resonance Imaging

For MRI scanning, induction was carried out with 5% isoflurane in medical air, followed by a maintenance dose of 1.5% isoflurane, which was finely tuned to ensure the animals remained stable during their time inside the magnet. The monitoring of the animals’ physiological status was facilitated by a sophisticated Small Animal Monitoring and Gating System capable of tracking respiration, body temperature, electrocardiogram, among other vitals. The respiration rate was constantly observed with a pressure-sensitive pad, aiming to keep it within 90-120 breaths per minute. To maintain the animals at a steady temperature of 37°C, warm air was circulated through the MRI scanner’s bore, with the temperature being verified by a rectal probe thermometer.

We used the 14T Magnex MRI scanner with Bruker Biospin AV Neo console equipped with a 1.2T/m gradient. We acquired high-resolution images using a Fast low angle shot magnetic resonance imaging (FLASH) sequence at two isotropic resolutions: 75 micrometers (TR: 100ms, TE: 2.08ms, Matrix: 192x128x192, BW: 50kHz with a total time of 24min22s) for broad anatomical context and 37.5 micrometers (TR: 100ms, TE: 4.4ms, Matrix: 384x256x384, BW: 50kHz, with a total time of 6h28min.) for ultra-high-resolution imaging, enabling an unprecedented level of detail in the visualization of the microvascular structure. This dual-resolution approach provides a comprehensive overview of the neurovascular unit’s architecture and function, crucial for understanding the underpinnings of neurophysiological and pathophysiological processes in the brain.

### Deep-Learning-Based Super-Resolution

We have provided a schematic drawing for the detailed steps in Fig 1. To perform the super-vised training, we first normalized the image of each scan. The 3D architecture for image super-resolution is a 3D convolutional neural network (CNN) including residual encoder and decoder layers^60,61,62^. It has a lightweight architecture with three resolution levels, each with a convolutional layer with 32 features and a Leaky rectified linear unit^63^. The CNN was trained to super-resolve images by a factor of 2 in every dimension, from 75μ isotropic to 37.5μm isotropic. Specifically, it receives as input a trilinear upsampled image (i.e., living on a 37.5μm space, but with smooth, 75μm-like features), and estimates a high-frequency residual that, when added to the input, produces a sharpened output. Training used the Adam optimizer^64^ to minimize the average absolute difference (i.e., L1 norm) between the target and predicted image intensities. Aggressive data augmentation was used during training, to mitigate the small size of the training dataset and improve generalization ability, including: noise and bias field injection; contrast, brightness, and gamma modification; and geometric augmentation including rotation, shearing, scaling, and nonlinear deformation^65,66^. The ex-vivo MRI scans from four mice were used for training and optimization. For each iteration, 1000 random local cubes with 48^3^ voxels were selected around the center of the brain MRI. The CNN was trained with 100 iterations. After training, the CNN was used to produce high-resolution images from the in vivo WT and AD mice. The trained CNN was used to produce high-resolution images for voxel-wise data analysis of MION-based CBV images of WT and AD mice.

**Figure 1.**
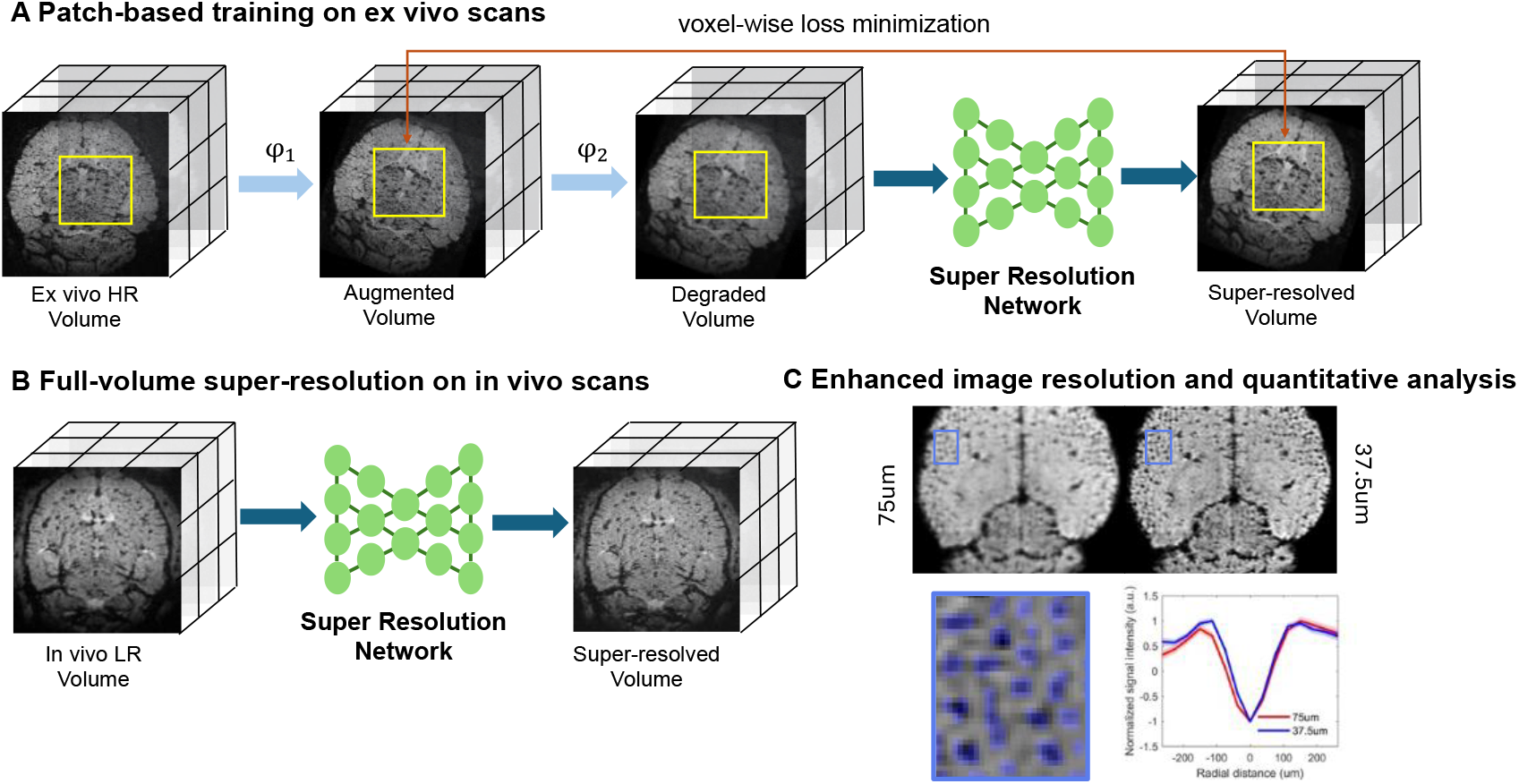
Overview of Super-Resolution network training and inference. (A) Patch-based training process on ex vivo scans. The workflow demonstrates the transformation from an ex vivo High-Resolution (HR) Volume to an augmented volume using spatial transforms, followed by the creation of a degraded volume with random noise and bias field injection; contrast, brightness, and gamma modification, which is then fed into the Super Resolution Network to generate a super-resolved volume. The voxel-wise loss minimization strategy is depicted as a feedback mechanism enhancing the network’s prediction. (B) Full-volume super-resolution on in vivo scans. This panel shows the application of the Super Resolution Network directly on an In vivo 75um Low-Resolution (LR) Volume to produce a 37.5um Super-resolved Volume. (C) Detailed visualization of super-resolution enhancement of axial 2D slices of CBV MRI images and line profiles of iron-induced T2* decay signals across selected micro-vessel ROIs, which was marked in blue color from the zoomed-in super-resolution-resolved images.

### MRI data processing

First, the gradient artifacts were removed using Analysis of Functional Neuroimages (AFNI) 3dUnifize function. Then, all of the high-resolution images were registered to the Australian Mouse Brain Mapping Consortium (AMBMC) template using AFNI 3dAllineate function. The brain mask made from the AMBMC atlas was applied to remove tissue out of brain. After removing the background signal obtained by Gaussian averaging, followed by sigmoid transform to convert and normalize the image, fast non-local mean filter to remove the noise, the MRI images were segmented for blood vessels based on signal intensity. The obtained binary vessel segmentation mask was averaged using a sliding window of size 450x450x450μm to obtain the vessel density map. Then a voxel wised t-test was applied to the WT and AD groups. The figures show the differential vessel density map of WT vs AD brain (p<0.01, FDR corrected).

### Statistics

To improve the stability and robustness of the differential map, the permutation method was applied. By randomly selecting 10 WT and 10 AD mice for two separate groups, we can create a permuted differential map PD. Significance is determined using a two-sample t-test (p<0.05). Each voxel of the permutated differential map can be calculated as:

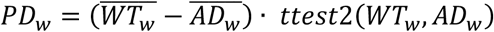

For the vessel density differential map, *WT*_*w*_ or *AD*_*w*_ represents the vessel density of a sliding window of 15x15x15 voxels, which was calculated by dividing the voxel numbers of segmented vessels by the voxel numbers of the brain mask. For the image intensity differential map, *WT*_*w*_ or *AD*_*w*_ represents the averaged z-score normalized image intensity within the brain mask of a sliding window of 15x15x15 voxels.

By randomly choosing the WT and AD mice 100 times, we created 100 permutated differential maps and resampled them back to their original size. A one-sample t-test (p<0.01) was then performed on these resampled permutated differential maps to generate the final differential map FD. Each voxel of the final differential map can be calculated as:

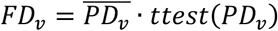

A similar permutation method was used to further confirm the significance of the final differential map. We randomly selected 5 WT and 5 AD mice as one group, and another 5 WT and 5 AD mice as the other group to create a control differential map. This control permutation was repeated 1000 times, and the distribution of the control differential maps in identified ROIs can be calculated and compared with the final differential map.

## Results

### Image processing overview based on the super-resolution resolved CBV images

We first implemented a super-resolution CNN network on the MION-based brain-wide CBV volumetric scans (Fig 1A), resulting in discernible enhancements in image resolution, from 75um to 37.5um isotopically (Fig 1B). There is a high localization fidelity of the vessels shown as the black dots with sharper edge in super-resolution resolved images when comparing with the original *in vivo* images acquired at 75um cubic resolution (Fig 1C). Meanwhile, we also analyzed the signal intensities of the super-resolution resolved and raw images, showing voxel-wise linear signal intensities changes across individual animals in both WT and AD groups (Supp Fig 1). Although no significant difference was detected between AD and WT mice in terms of the fitted linear function parameters (Supp Fig 1B), in the following studies, the voxel-wise analysis for size-dependent vascular compartments was based on the segmented micro-vascular masks to calculate the VD, but not on the signal intensity between the two groups of animals. It should be noted that the MION-based capillary density estimates from the parenchyma voxels were processed from the raw images after segmenting out the discernible micro-vessels based on the super-resolution-resolved images.

### Micro-vascular segmentation from super resolution MRI Data

As described in the data pre-processing pipeline (Fig 2A, details in the Method section), normalized high resolution images were inverted so that the vessel voxels showed high signal intensity, which was presented in the histogram plot of the 3D CBV image datasets (Fig 2B). Given the T2*-weighted amplification effect, the micro-vessels can be well identified from the MION-based CBV MRI images, which is shown in the 3D segmented cerebrovasulature (Fig 2C). By setting the threshold from top 2% to 50% of the total voxels from the histogram, the segmented 3D cerebrovasculature rendering view and 2D masks was presented in Fig 2D. These progressive stages of vessel segmentation revealed the hierarchical organization of the cerebrovascular network that become increasingly intricate at higher threshold levels. Furthermore, we examined the distribution of vessel size in both WT and AD mice, focusing on a range of volumetric fraction thresholds (2-50%), which serve as proxies for vessel size - where larger thresholds correspond to smaller vessels. Mean normalized signal intensity profiles of vessels identified at different thresholds were plotted to present the spatial distribution of micro-vessels in voxel-wise formality, showing progressively decreased spatial coverage of identified vessels when the threshold was increased from 2% to 50% in both groups (Fig 3A,C) Meanwhile, The full-width-of-half-maximum (FWHM) for vessels identified at different thresholds were presented in the violin plots, highlighting the comparable mean and median vessel sizes between two groups, as well as the distribution of vessel at different sizes for each threshold (Fig 3B,D). Statistical analysis showed no significant differences in vessel size comparison between WT and AD models across varied thresholds. (Fig 3E), suggesting the iron-induced T2*-decay presented similar global amplification effects between two groups. It further supported the brain-wide VD mapping to elucidate the region-specific alterations in degenerated AD brains.

**Figure 2:**
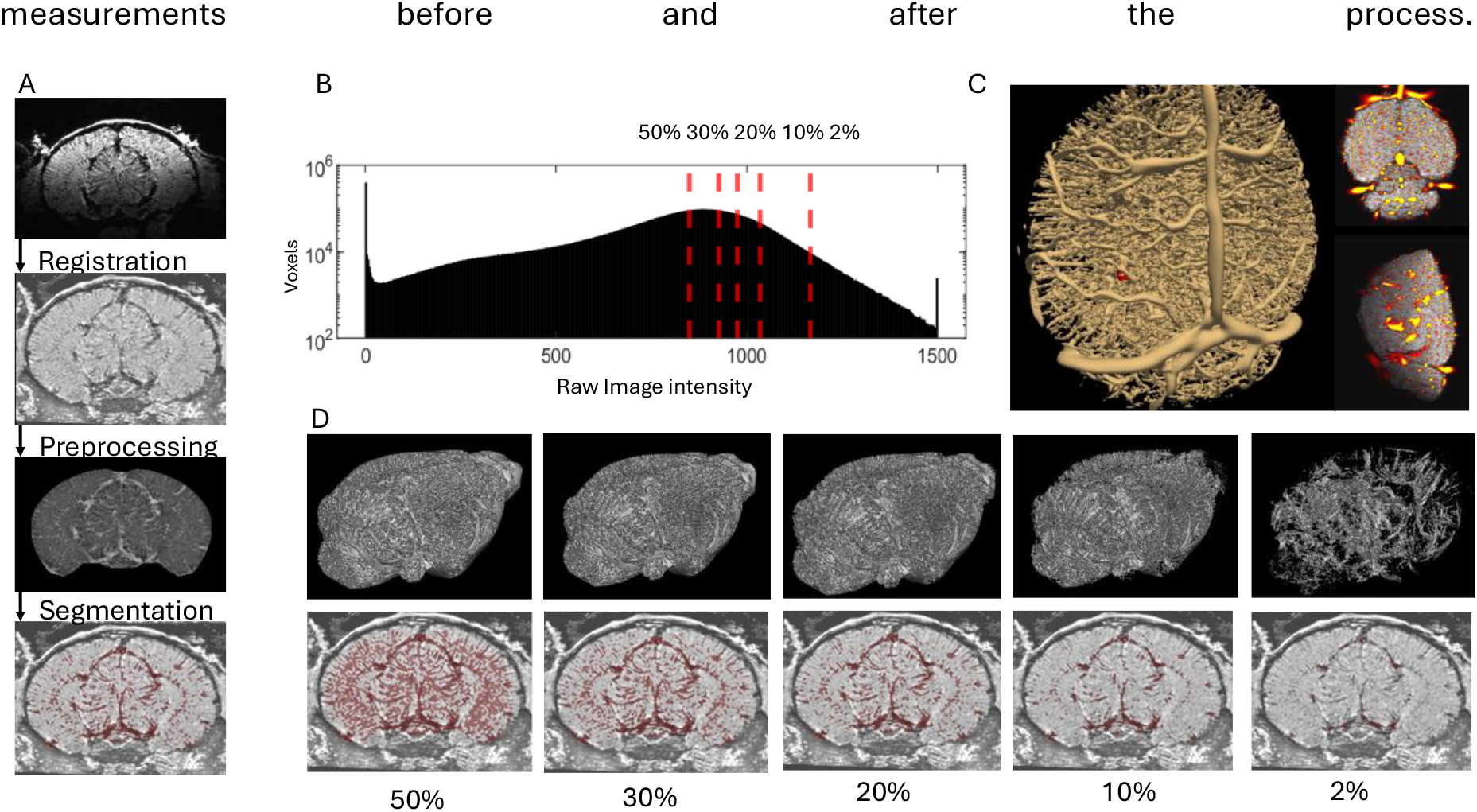
Assessment of vessel segmentation. (A) Pipeline of data processing. (B) The histogram of voxel intensity distribution, with a threshold range marked by red dashed lines to delineate the segmentation parameters for vascular structures. (C) The representative 3D rendering view of segmented micro-vessels and overlaid micro vessels on CBV MRI images. (D) Visualization of vessel segmentation at varying threshold. The top row provides the 3D rendering of the vascular structure; The bottom row provides a fusion of the structural and vascular images, with the vasculature superimposed in red over the grayscale anatomical slices.

**Figure 3:**
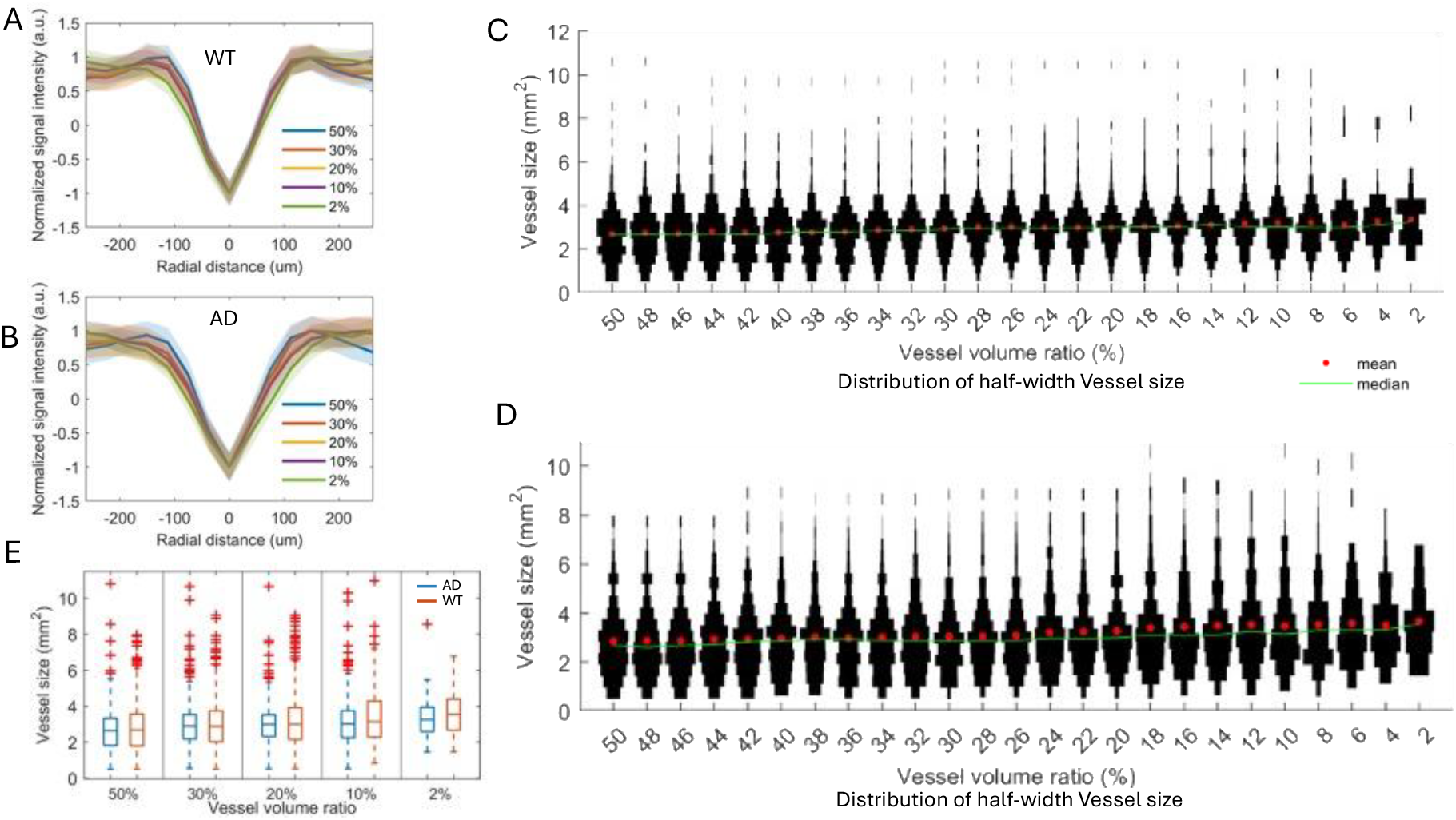
Quantitative Analysis of Vascular Volume Fractions. (A&B) The normalized average distribution of voxel intensity within specified volumetric fraction thresholds, ranging from 2% to 50%, with color bands representing confidence intervals, in wildtype (A) and AD mouse groups (B). (C&D) The violin plot distribution of voxel intensities at varying volumetric fraction thresholds, with the mean and median values highlighted, in wild type model (C) and AD model (D). (E) The comparison between WT and AD models, showing the vessel volume ratio across different thresholds. No significant difference was detected between two groups.

### Brain-wide vessel size-dependent alteration of VD in AD brains

We first identified VD maps, i.e. the number of vessel voxels of a given kernel of 15x15x15 voxels, based on the segmented micro-vessels with a threshold of 50% of the pre-processed CBV-based MRI images. Fig 4 showed the differential VD maps between AD and WT mice from different coronals slices, highlighting altered VD in the AD mouse brains. No significant difference was detected in majority of the cortex and a large portion of the hippocampus. Nevertheless, several key brain areas involved in the cognitive decline during AD brain degeneration showed vascular alterations. The anterior cingulate cortex (ACC) and medial entorhinal cortex (mEC) presented decreased VD in contrast to the increased VD observed in the Piriform area (Pir), anterior insular cortex (AI), and lateral entorhinal cortex (lEC). The ventral part of hippocampus, including dentate gyrus (DG) and amydalohippocampal areas, also showed increased VD. Increased VD was also observed in the medial septal area (MS), central and geniculate thalamic nuclei, and ventral tegmental area (VTA). At the midbrain, increased VD was observed in the inferior collicus (IC), pedunculopontine tegmental nucleus (PPT), laterodorsal tegmental nucleus (LDT) and periaqueductal gray (PAG). Fig 4B showed the statistical comparison of the VD identified from the selected region-of-interests (ROIs).

**Figure 4:**
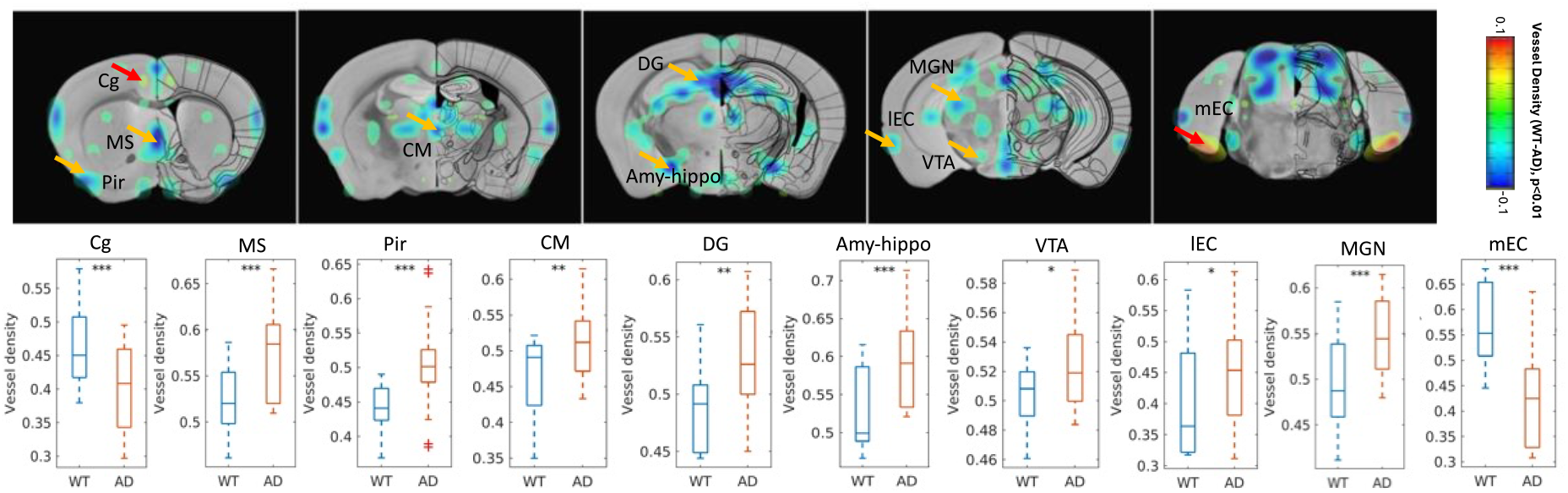
Vessel density difference between WT and AD mice. Statistical parametric map to show the VD difference between WT and AD mice based on the micro vessels characterized with 50% threshold (two-tailed t-test, FDR P ≤ 0.01, permutated for 1000 times, color-coded scale of vessel density difference). Differential maps were overlaid with the mouse brain atlas (the 2D coronal slices were shown different brain regions from rostral to caudal axis). The bar plots to show the ROI-specific mean vascular density from WT and AD mice (N_AD_ = 12, N_WT_ = 11, *p<0.05, **p<0.01, ***p<0.001)

To elucidate the refined alteration specific to vascular compartments at different vessel sizes in AD brains, we produced the VD differential maps at thresholds:2%, 2-10%, 10-20%, 20-30%, 30-40%, and 40-50% (Fig 5). Consistent with the accumulative VD differential analysis, no significant difference was observed in majority of the cortex and the hippocampus. However, the decreased VD in mEC was mainly observed at the threshold of 2-10% threshold and increased VD in Pir, AI, and lEC was detected at the threshold of 2-30%. Variable VD differences were observed near ventricle areas, showing deceased VD at the thresholds of 2% and 30-50% but increased VD at the threshold of 2-30%. Interestingly, the increased VD was also observed at the pontine area at the threshold of 30-50% but not observed in the accumulative VD differential maps. These results suggested that VD alteration in the AD brains showed both size- and region-specific patterns, which can be well characterized by the MION-based CBV MRI.

**Figure 5:**
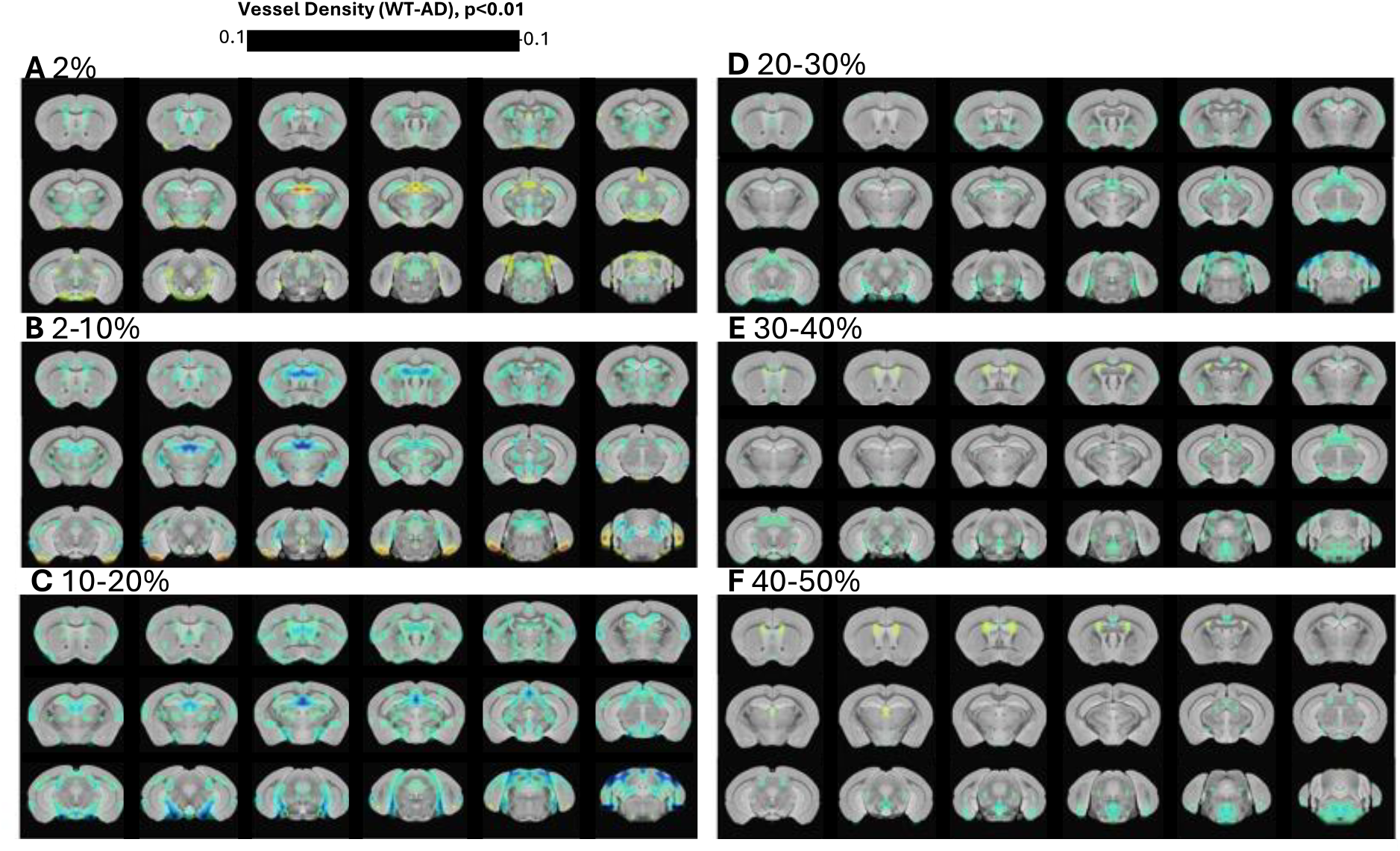
Vessel density difference at different thresholds between WT and AD mice. Statistical parametric map to show the VD difference between WT and AD mice based on the micro vessels with different vascular segmentation thresholds from 2% to 50%(A-E) (two-tailed t-test, FDR P ≤ 0.01, permutated for 1000 times, N_AD_ = 12, N_WT_ = 11).

### Brain-wide capillary VD alteration in AD brains

Although MION-based CBV MRI enabled the segmentation of micro-vessels with tens of micron diameter based on the iron-induced T2* amplification effect, it remains impossible to dissect the capillaries given its limited spatial resolution. Nevertheless, the T2*-weighted signal intensity changes of parenchyma voxels provided an alternative scheme to estimate the capillary VD. Fig 6 presented the capillary VD differential maps between WT and AD mice from parenchyma voxels of the CBV-based images by excluding the segmented micro-vessels based on the 50% threshold. Since the VD is estimated based on the CBV-weighted signal intensity, the positive signs of the differential maps indicated the increased capillary VD in AD mice. Interestingly, increased capillary VD was observed in the somatosensory, visual, retrosplenial cortices, Pir, and lEC of AD mice, but ACC showed decreased capillary VD. Whereas, dramatically decreased capillary VD was observed in the subcortical areas, including the thalamus, hippocampus, hypothalamus midbrain and pontine areas. Fig 6B highlighted the ROI-based quantitative analysis of capillary VD in the hippocampus and somatosensory cortices, showing significantly altered VD in AD brains. This non-invasive vascular mapping scheme has enabled brain-wide characterization of the capillary density alteration as AD-related vascular anomalies. To be noted, we used permutation analysis to enhance the reliability of test results. By randomly shuffling data and recalculating test statistics across 1000 permutations, this approach generates a distribution of outcomes under the null hypothesis, allowing for a more accurate estimation of group analysis results(Supp Fig 2 and 3).

**Figure 6:**
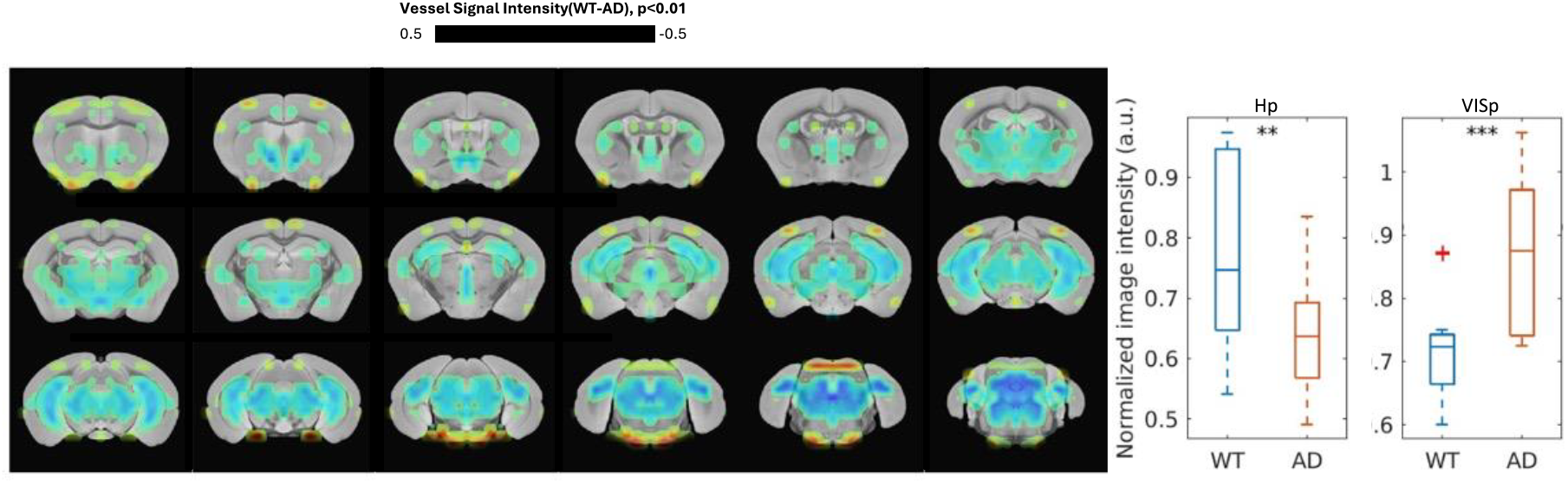
Capillary density difference between WT and AD mice. Statistical parametric map to show the capillary density difference between WT and AD mice by deducting micro vessels detected (threshold 50%). (two-tailed t-test, FDR P ≤ 0.01) The bar plots to show the ROI-specific mean capillary density of Hippocampus and somatosensory cortex from WT and AD mice(N_AD_ = 12, N_WT_ = 11, *p<0.05, **p<0.01, ***p<0.001)

## Discussion

This study extends the capacity of MION-based CBV MRI for brain-wide cerebrovascular density mapping, enabling the detection of cerebrovascular alterations in degenerated AD brains. There were two ongoing challenges for the MRI-based micro-vascular mapping. One is the intrinsic limitation of the spatial resolution of proton-based *in vivo* MRI. The other is the appropriate MR contrast that can be used to reliably identify micro-vessels aligning with various orientations across the brain. This work implemented the deep learning-based super-resolution approach to map the amplified micro-vessels from T2*-weighted images by injecting the iron oxide nanoparticles into the blood.

To achieve the ultra-high spatial resolution for *in vivo* mouse brain MRI scanning, we applied the 14T MR scanner with an implantable RF coil to boost the SNR^67^. Although the ultra-high field MRI has the capability to perform 15um isotropic resolution ex vivo scanning ^68,69^. this MION-based MRI would not gain the similar resolution for several reasons: i. the iron particle-induced faster T2* decay leads to much lower SNR of the brain images; ii. since ultra-high field MRI shows high sensitivity to field inhomogeneity, the high bandwidth for frequency encoding is needed to ensure short TE to avoid large field distortion, which would also limit the SNR and maximal matrix size for high resolution scanning; iii. the *in vivo* scanning scheme also limit the extended acquisition time which is usually needed to obtain sufficient SNR at the isotropic resolution of 10-20 μm. In this work, we have applied high order shimming to ensure a ∼60-75Hz at 50% bandwidth for the *in vivo* whole brain field homogeneity with iron injected into blood at 14T, i.e. 600MHz, which is less than 0.1ppm. It should be noted that the implanted RF coil over the mouse head reduced the air-tissue interface artifacts and ensured this better field homogeneity when acquiring the ultra-high resolution MION-based MRI^67^.

We have implemented the deep learning-based super resolution scheme to improve the CBV MRI image from 75μm to 37.5μm isotropic resolution. Our results have shown high spatial fidelity of the micro-vessels highlighted by MION contrast in the T2*-weighted images (Fig 1C), enabling refined segmentation of the vessels to different compartments. Besides the resolved higher resolution, this method could alter the signal intensity of the images. Although there were proportional changes of the signal intensity of brain voxels as shown in Supp Fig 1, specific caution should be given when analyzing the super resolution-resolved dataset based on the signal intensity between groups. Such a change might be caused by the different signal intensity voxel-wise distribution between *in vivo* and *ex vivo* scans, considering that the model was trained on *ex vivo* scans and tested on in-vivo ones. The inherent acquisition difference - namely domain shift, is a common issue in existing super-resolution methods^70,71^. Also, previous application of super-resolution approaches to improve the low-field MRI images have demonstrated the high fidelity for morphometrical segmentation studies^72,73^. Therefore, we applied the vascular density maps only based on the segmented vascular masks and the capillary density comparison was performed from the raw dataset after segmenting the larger micro-vessels.

Other preclinical brain-wide vascular mapping methods, e.g. doppler-based ultrasound or optoacoustic imaging, either relies on the flow or the hemoglobin contents^74-76^. These methods provided single digit micron spatial resolution to map brain-wide cerebrovasculature, but spectral penetration issues and vascular orientation lead to less detectability of micro-vessels located in the deeper brain areas or vessels aligned perpendicular to the doppler beams. In contrast, despite the lack of spatial resolution of MRI, the iron-induced T2* effect from blood could amplify the micro-vessels with diameters of 20μm or above to be well characterized in CBV MRI images ^77,78^. The typical size of penetrating micro-vessels identified with high resolution MRI is characterized by comparing mean vascular distance between nearest arterioles and venules detected by optical imaging methods^78-81^. Due to the T2*-based extravascular effect, the line-profiles of decayed signals across micro-vessels showed the FWHM near 100 μm, which also explained the much larger thresholds of the total voxels when segmenting the micro-vessels than the typical 2-4% partial volume fraction of cerebrovasculature^82-84^. Despite the over-estimated volume fraction of segmented micro-vessels from CBV-based MR images, the typical vessel size distribution between AD and WT mice are comparable from the super-resolution resolved images (Fig 3), enabling the voxel-wised group analysis to measure the region-specific micro-vessel density changes from the degenerated AD brains. Furthermore, by excluding the identifiable micro-vessels, the iron-induced signal intensity reduction from parenchyma voxels would indicate the capillary density given its low leakage rate through the blood-brain-barrier, which has been previously applied to map the tumor vascularization in rodent brains^85^. This vascular imaging scheme based on iron-induced T2*-weighted signal reduction has been applied to estimate dynamic vascular density changes in APP32 transgenic AD mice ^20^. This work has applied 1mm slice thickness and estimating vessel size based on modeling the iron-induced susceptibility changes from T2/T2* weighted images acquired before and after iron injection with apparent diffusion coefficient (ADC) estimates^86,87^. Different from our work, this previous study cannot differentiate the different vascular compartment contribution and the altered VD changes can be confounded by altered brain-wide T2/T2*-weighting patterns before the iron injection, as well as the region-specific ADC changes of AD mice.

Our work offers an advanced mapping scheme to compare brain-wide vascular density between AD and WT animals based on the separated vascular compartments. We have demonstrated the 3D vasculature distribution with 50% volume thresholds, covering the majority micro-vessels identifiable with the MION-based CBV MRI (Fig 2C, D). Also, by adjusting the thresholds, we can estimate the vessel size-dependent VD differential maps to further identify vascular alteration in degenerated AD brains. This unique mapping scheme is well exemplified when pinpoint affected brains regions showing specific differential VD features. For example, decreased VD at mEC and increased VD at lEC of AD mice were first identified in the 50% threshold-based differential maps, which could be further specified to vessel size-specific differential maps with 2-10% threshold. In contrast, the deceased VD in ACC was only identified in the 50% threshold accumulative differential map, and increased VD in the pontine area was only identified in the vessel size-specific differential maps with 30-50% threshold but not detectable in the 50% threshold maps. These vessel size-dependent VD difference is further assessed by the capillary VD differential maps, showing increased VD in a few cortical areas, but decreased VD in majority subcortical areas. It should be noted that these highly region-specific VD changes with varied signs in degenerated AD brains further eliminate the group variance effects due to different iron administration and clearance between AD and WT mice.

This vascular compartment-specific VD differential maps could help elucidate the confounding observations of VD alteration reported in AD patients’ brain specimens and AD-like rodent models. We have observed that hippocampus showed increased VD in the DG area for micro-vessels identified with 50% threshold, which may be relevant to reported hippocampal VD increase in human brain specimens^88-90^. Whereas, the capillary VD of hippocampus is significantly decreased across the hippocampal areas, presenting consistent observation with other reported human studies ^25,26,91^. Brain-wide VD reduction, in particular, the subcortical areas including hippocampus, thalamus, and basal ganglion, has been reported in multiple APP/PS1 and Tg transgenic mouse lines^9-189,92^.

Interestingly, increased cortical VD was reported in the Tau(P30IL) transgenic mice and several human specimen studies^4,93,94^. We also observed the increased capillary VD in multiple cortical areas of the APP/PS1 mouse line, which was previously reported to show no change or decreased VD by two-photon imaging or immunostaining^18,95^. And, no significant cortical VD changes were observed based on the 50% threshold, which may reconcile our observation with the previous APP/PS1 studies. Whereas the decreased VD was only observed in the ACC area in both capillary and micro-vessel (50%) VD differential maps, demonstrating its essential role in AD pathophysiology.

This brain-wide vascular compartment-specific VD mapping enables the characterization of vascular alterations in degenerated AD mouse brains. Different from previous CBV MRI studies for vascular size imaging, we applied super-resolution approach to assist the direct segmentation of the micro-vessels from *in vivo* high-resolution MRI images. Several limitations should be considered when implementing this method for quantitative analysis of VD. First, the iron-induced T2* amplification effect relies on the dosage of iron particles injected into blood. It is critical to compare global iron-induced T2* decay among animals of different groups to avoid the statistic errors caused by group variance. Second, super-resolution approaches model inferring details to fit its learned data distribution, which could result from overfitting or biases in the training ex-vivo dataset. We will further optimize the machine learning neural network by training the ultra-high resolution *in vivo* images in the future study with robust validation and domain-adaptive refinement of the model ^96^. Thirdly, although the vascular compartments can be categorized based on varied thresholds when segmenting micro vessels from CBV MRI image, the characterization criteria of the vessel size is arbitrary, only showing proportional relevance to the true vessel sizes. Future studies should be focused on implementing the vessel size modeling scheme with ultra-high resolution MRI to characterize vascular density with higher accuracy. Our work has shown the transformative potential of ultra-high resolution CBV MRI to identify cerebrovascular alterations associated with AD, paving the way for improved diagnostic capabilities and therapeutic interventions.

---

**Supplemental Figure 1:**
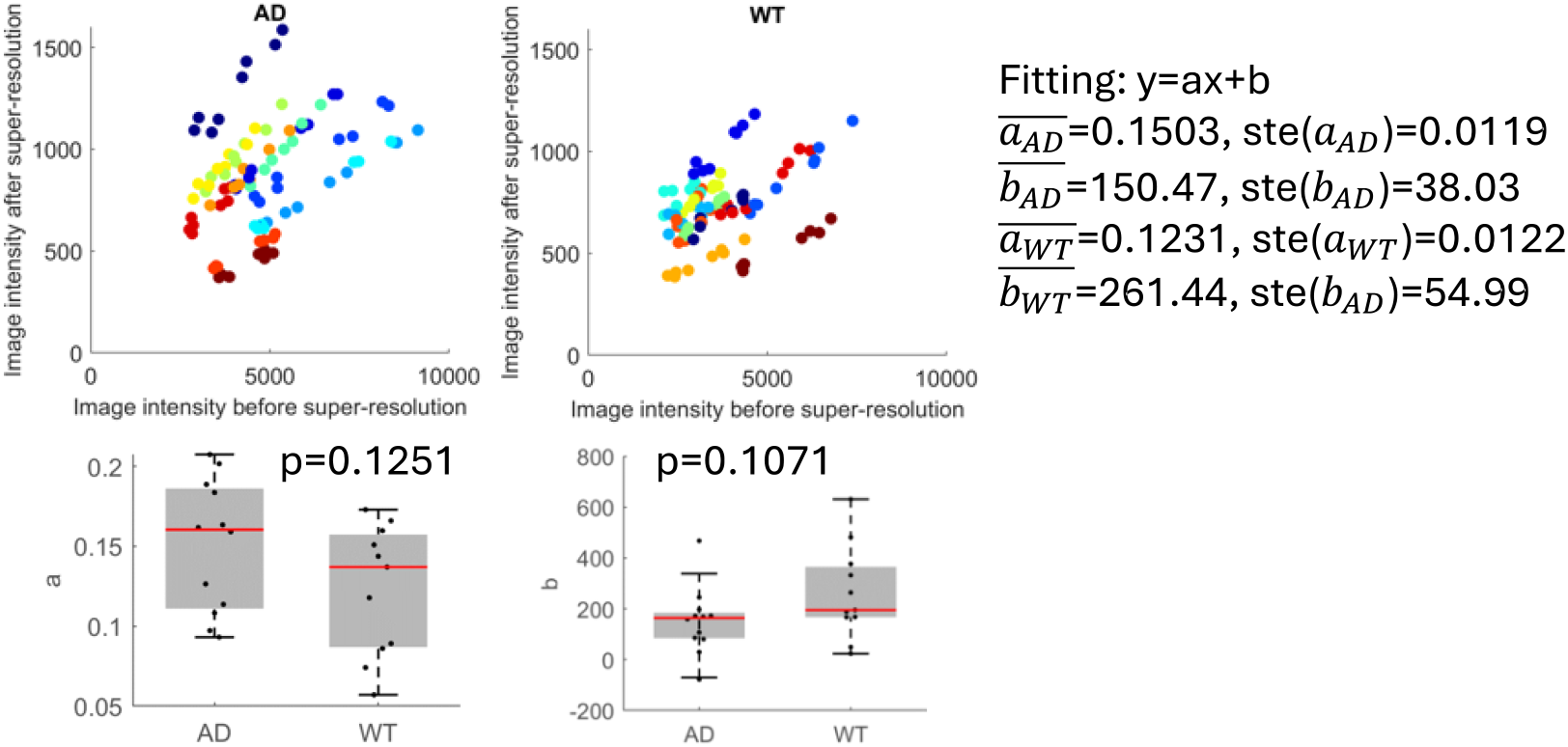
Comparison of Image Intensity Enhancement and Distribution Analysis in AD and WT Mice Pre- and Post-Super-Resolution Processing. (A&B) Scatter plot displaying the correlation between image intensity values before and after super-resolution in AD mouse (A) and WT mouse (B) brain images. Each point represents a specific region within the brain, color-coded to indicate the relative location across multiple samples, demonstrating the enhancement of image intensity through super-resolution techniques. (C) Boxplot comparison of the parameter ‘a’ derived from the super-resolution process between AD and WT mice. ‘a’ represents a metric of image quality or specific feature enhancement. (D) Boxplot comparison of another parameter ‘b’, also derived from the super-resolution process, between AD and WT groups. This metric is related to image contrast and noise characteristics. There is no significant difference in neither ‘a’ or ‘b’ between AD and WT groups. The scatter plots (A and B) illustrate the effectiveness of super-resolution in enhancing image intensities across both AD and WT models, while the boxplots (C and D) provide a statistical measure of the variation in specific image quality parameters resulting from super-resolution, offering insights into the differential effects of the process on pathological versus normal brain tissue.

**Supplemental Figure 2.**
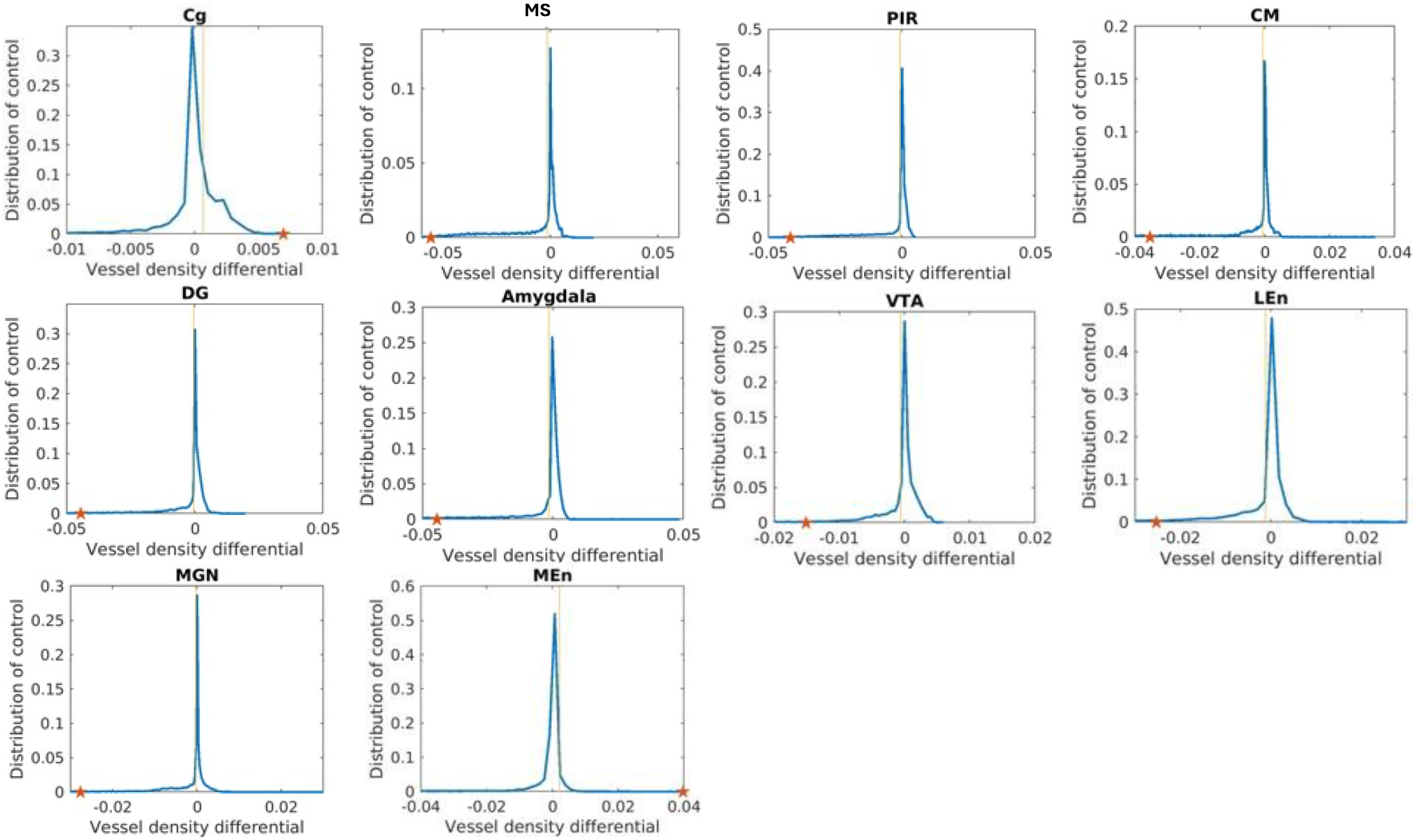
Permutation control results to show the histogram distribution of vascular density difference from selected ROIs based on randomized animals from two groups.

**Supplemental Figure 3.**
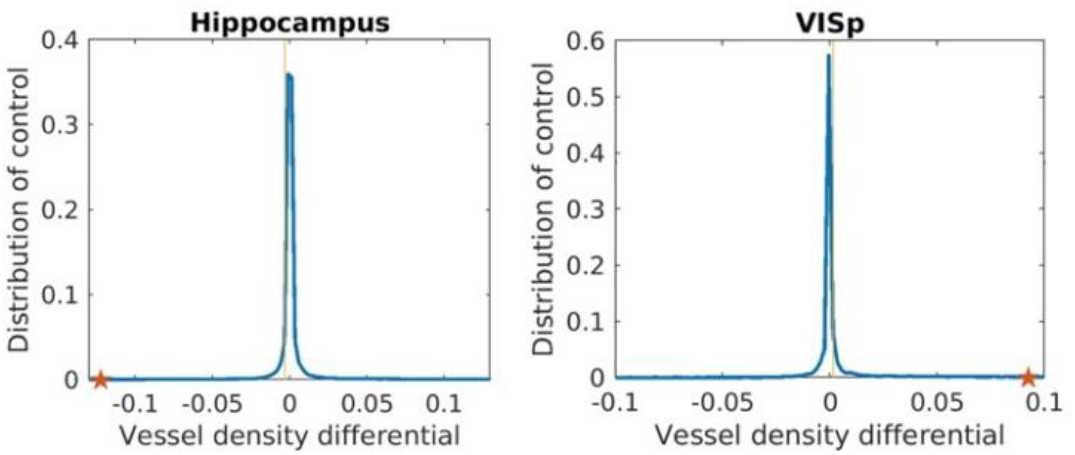
Permutation control results to show the histogram distribution of capillary density difference from selected ROIs based on randomized animals from two groups.

